# Reframing dopamine: Controlling the controller at the limbic-motor interface

**DOI:** 10.1101/2023.03.25.534180

**Authors:** Kevin Lloyd, Peter Dayan

## Abstract

Pavlovian influences notoriously interfere with operant behaviour; various such may be associated with the release of the neuromodulator dopamine in the nucleus accumbens. One role for cognitive control is suppressing these influences. Here, using the examples of active avoidance and omission behaviour, we examine the possibility that one instrument of control is direct manipulation of the dopamine signal itself.

---

It can be difficult for animals to learn to act vigorously to avoid predicted punishment, or to withhold actions to gain potential reward (Crockett, Clark, & Robbins, 2009; Dayan, Niv, Seymour, & Daw, 2006; Guitart-Masip, Beierholm, Dolan, Duzel, & Dayan, 2011; Swart et al., 2017). One cause of this may be the neuromodulator dopamine (DA), in a conflict between its dual roles in positive reinforcement and motivational vigour (Collins & Frank, 2014; Salamone & Correa, 2012; Schultz, Dayan, & Montague, 1997). Evidence from canonical versions of these paradigms, which appear to inspire Pavlovian-instrumental conflict, suggests that DA in the core of the nucleus accumbens (NAc) follows its motivational, rather than its reinforcement, role (Gentry, Lee, & Roesch, 2016; Oleson & Cheer, 2013; Syed et al., 2016). Here, we build on a model (Lloyd & Dayan, 2019) of an heuristic suggestion (Boureau & Dayan, 2011) for active avoidance, characterizing DA dynamics in NAc as resulting from internal cognitive control actions.

Gentry et al. (2016) used fast-scan cyclic voltammetry (FSCV) to examine DA release in the core of NAc during performance of a mixed-valence task. In one class of trials, rats heard a tone telling them that they had to press a lever within a 10 s response window to avoid a shock (**Fig. 1a**). Half the animals often struggled to respond actively in time (**Fig. 1b**,**c**), and showed higher rates of freezing — a typical Pavlovian-instrumental conflict. However, across the population, on trials when they did press, cue-elicited DA release was similar just after shock or reward cues (**Fig. 1d**), notably being stimulated rather than suppressed. This finding has deeper roots in the literature (Brischoux, Chakraborty, Brierley, & Ungless, 2009; Budygin et al., 2012; de Jong et al., 2019; Oleson & Cheer, 2013; Young, 2004).

**Figure 1:**
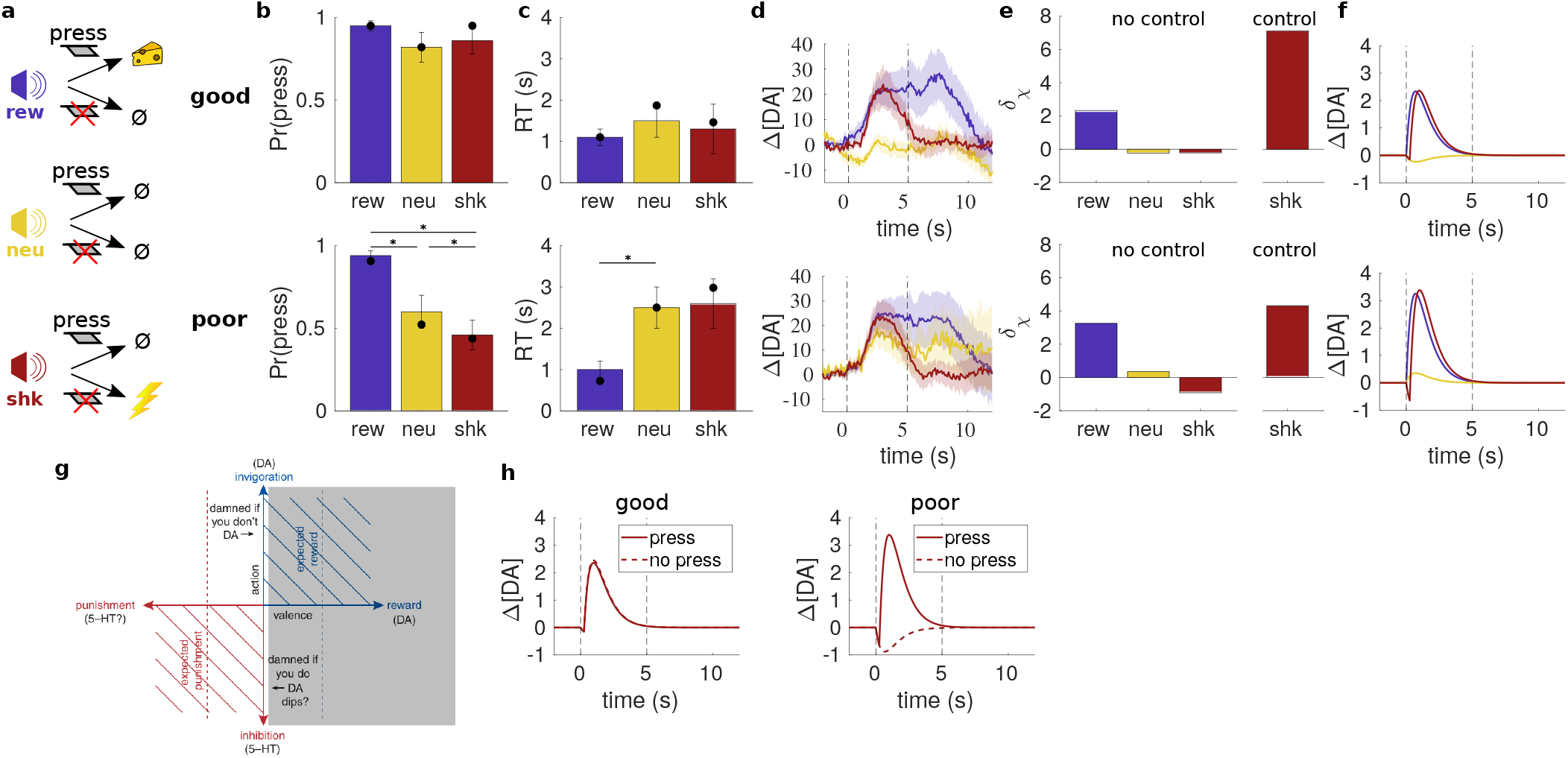
Mixed-valence task (Gentry et al., 2016). (**a**) On each trial, a tone indicated whether a lever press within 10 s would yield reward, have no consequence, or avoid a scheduled shock. (**b**,**c**) Good-avoiders (upper) pressed often and quickly; poor-avoiders (lower) pressed less often and more slowly on shock and neutral trials (^*^indicates significance, *p <* .05). Filled circles indicate model fit. (**d**) Average cue-aligned NAc DA release (±SEM) on press trials for each trial type (vertical dashed lines indicate cue onset at 0 s and lever insertion at 5 s). (**e**) Cue-evoked TD errors predicted by model. (**f**) Average DA release predicted by model on press trials. (**g**) Putative shifting of origin leftwards in the affective circumplex to promote DA release and approach to safety in the active avoidance case (adapted from Boureau & Dayan, 2011). (**h**) Predicted DA release for press vs. no-press shock trials. (Figures b–d adapted from Gentry et al., 2016).

Lloyd and Dayan (2019) suggested this DA signal was a partial temporal difference (TD) prediction error that reflects the positive relative utility of avoiding the shock, i.e., the value of safety. However, they did not provide a process account of the reframing required to turn an at best negative situation into one that appears positive. Here, drawing also on the mirrored results about DA dynamics in the core of NAc when actions must be withheld to gain rewards (Syed et al., 2016), we suggest that cognitive control, along with other means of influencing action and inaction (Cain, 2019), exploits the effects of DA to help overcome Pavlovianinstrumental conflict.

In Syed et al. (2016), one of four auditory cues indicated whether rats had to leave a nosepoke (‘Go’) and execute an active response (press a lever twice) or stay in the nose-poke until the tone turned off (‘No-Go’) in order to get a small or large reward (**Fig. 2a**). Animals were reliably successful on Go large-reward (GL) trials, but less so on No-Go large-reward (NGL) trials (**Fig. 2b**). We attribute this to Pavlovian misbehaviour caused by the prospect of a large reward, consistent with the faster ultimate reaction time on successful large-reward trials in both Go and No-Go conditions (**Fig. 2c**). Mirroring the case of active avoidance (Gentry et al., 2016), on successful NGL trials, after a minor peak, there was a suppression of DA below baseline during the No-Go period (followed by a rise at movement initiation), despite the prospect of large reward (**Fig. 2d**); by contrast, on successful GL trials, there was a marked early increase.

**Figure 2:**
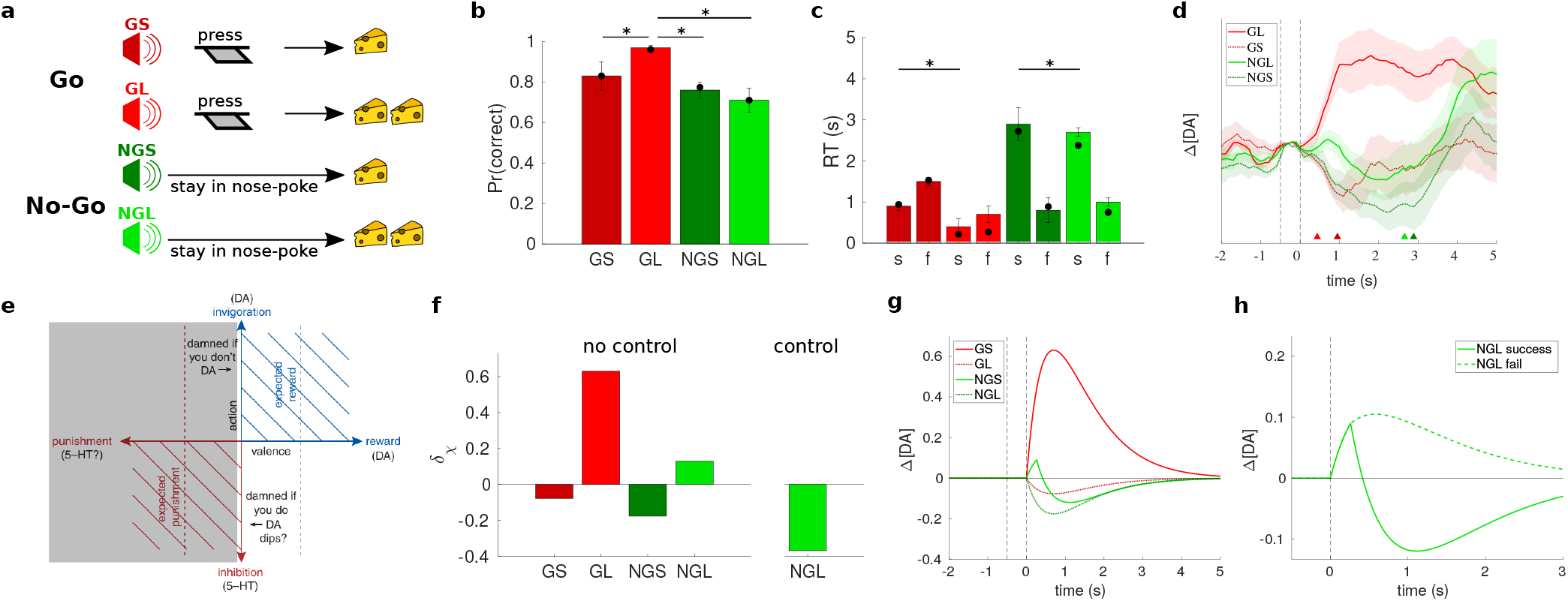
Go/No-Go task (Syed et al., 2016). (**a**) On each trial, a tone indicated whether the animal should leave the nose-poke and press a lever (‘Go’) to gain a small (GS) or large (GL) reward; or remain in the nose-poke until the tone turns off (‘No-Go’) to gain a small (NGS) or large (NGL) reward. (**b**,**c**) Average success rates and RTs (±SEM) for each trial type; the latter are split further into successful (s) and failed (f) trials (^*^indicates significance, *p <* .05). Filled circles indicate model fit. (**d**) Average cue-aligned change in DA (±SEM) for each trial type on successful trials; triangles indicate mean RTs. (**e**) Putative shifting of origin rightwards in the affective circumplex to suppress action in light of predicted reward (adapted from Boureau & Dayan, 2011). (**f**) Cue-evoked TD errors predicted by model. (**g**) Average DA predicted by model on success trials. (**h**) Average DA predicted by model on successful vs. failed NGL trials. (Figures b–d adapted from Syed et al., 2016.)

We formalize the problems, and the suggested cognitive control solution, by extending a conventional model of Pavlovian effects on behaviour (Huys et al., 2011). This adds *ωV* (*s*) to the propensity to emit an action at state *s*, where *V* (*s*) is the current estimate of the state’s long-run value and *ω* is a parameter. Assuming *ω >* 0, then if *V* (*s*) *>* 0 (e.g., from somewhat successful NGL performance), this factor *boosts* the propensity to act. Noting (a) that the TD error (Sutton, 1988) typically associated with state *s*_*t*+1_,

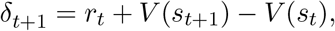

is just the *value* of that state, *V* (*s*_*t*+1_), if there is no extrinsic reward at that moment (*r*_*t*_ = 0) and no precise prior expectation (*V* (*s*_*t*_) = 0); and (b) the dopaminergic realization of this TD error (Kim et al., 2020; Montague, Dayan, & Sejnowski, 1996; Schultz et al., 1997; Starkweather & Uchida, 2021), a possible realization of this Pavlovian effect is the incentive salience-associated release of DA (Berridge, 2007), energizing action (McClure, Daw, & Montague, 2003), perhaps via its action on direct and indirect pathways in the striatum (Collins & Frank, 2014).

Conversely, if *V* (*s*) *<* 0 (e.g., from partially incompetent active avoidance), the Pavlovian factor, *ωV* (*s*), *suppresses* the propensity to act, e.g., by promoting freezing or withdrawal. It is less clear that this arises from below-baseline DA at *V* (*s*_*t*+1_) *<* 0; indeed, this condition has been associated with the activity of a separate system that is opponent to DA (Boureau & Dayan, 2011; Daw, Kakade, & Dayan, 2002).

For convenience, write *s*_pre_ for the state before the disambiguating cue in either experiment, with *V* (*s*_pre_)≃ 0 (from long, subjectively uncertain, inter-trial intervals), and *s*_*χ*_, *χ* ∈ {shk, ngl}for the states of interest inspired by the cue. Then, for shock trials in the mixed-valence task and imperfect avoidance, *V* (*s*_shk_) *<* 0. Thus, putatively,

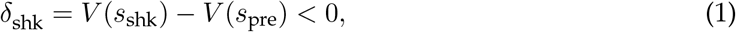

promoting Pavlovian inhibition which, as positive feedback, would make avoidance harder, thus making *V* (*s*_shk_) more negative and exacerbating the problem. For the immediate response to the NGL cue in the Go/No-Go task, we have the opposite problem. Partial success at No-Go, and thus large rewards, would make *V* (*s*_ngl_) *>* 0, with

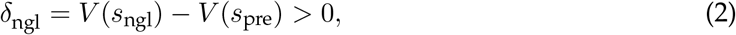

promoting Pavlovian action, No-Go failure, and ultimately a decrease in *V* (*s*_ngl_), lessening the misbehaviour. This slow negative feedback could even lead to oscillations.

Our central conceit is that the coupling of DA with action provides both the opportunity and need for cognitive control in which DA release is manipulated by a *reframing* of values. This generalizes the suggestion (Boureau & Dayan, 2011; Lloyd & Dayan, 2019) that the origin of the valence axis of the affective circumplex can be adjusted.

Control might just determine a particular trajectory for DA release. We explore a more limited construct in which it induces new, counterfactual (Bennett, Davidson, & Niv, 2022), states — in the simplest case, substituting for *s*_pre_ — associated with default expectations and potentially, though unnecessary here, with default actions. This induction then influences DA. For active avoidance, cognitive control instills a state *s*_fail_, with *V* (*s*_fail_) ≪ 0 quantifying the full explicit cost of the shock. Then,

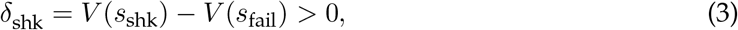

promoting Pavlovian action. This relocation of the origin of the affective circumplex to the negative affective value associated with presumed failure and thus the shock (**Fig. 1g**) harmonizes Pavlovian and instrumental control in the service of active responding — and would explain the positive (“damned if you don’t”) DA transient for successful avoidance in Gentry et al. (2016).

To test this, we fitted a model (see Methods) that incorporates Pavlovian influences, via an effective value of *ω*, and a probability of employing control to the animals’ behaviour (**Fig. 1b**,**c**). Averaging over the resulting mixture of differential dopaminergic TD errors for no-control vs. control shock trials (**Fig. 1e**) then indeed implies a net-positive DA signal on trials where animals successfully avoid shock (**Fig. 1f)**. The predicted suppression of DA release for poor avoiders on failed avoidance trials (**Fig. 1h**) would be consistent with such failures of control, and with observations (Oleson, Gentry, Chioma, & Cheer, 2012; Wenzel et al., 2018) that enhanced or suppressed DA release given a warning cue predicts successful or failed active avoidance. Of course, given small *ω*, successful avoidance could be achieved without control.

For NGL trials in the Go/No-Go paradigm, we similarly consider cognitive control as instilling a state *s*_succ_ with *V* (*s*_succ_) ≫ 0 quantifying the full value of succeeding in the No-Go requirement. Then

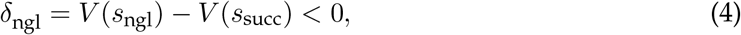

again harmonizing Pavlovian and instrumental control by facilitating inaction. This amounts to moving the origin of the affective circumplex to the positive value associated with presumed success in NGL (**Fig. 2e**), switching the sign of the TD error (**Fig. 2f**) and leading to suppression of DA release. The brief initial increase on successful NGL trials (**Fig. 2g**) might arise before control is exerted, something we would need a finer timescale model to examine. Here, control failure, associated with failed NGL trials, should lead to enhanced DA release (**Fig. 2h**). Apparent trends in this direction were not, however, found to be significant (**Fig. S1**), though the relatively small sample size and large variability in the voltammetry signal may obscure such differences.

An important remaining problem with the proposed reframing is the apparent absence of learning. For instance, if the DA signal of equation 3 is positive, why does normal plasticity, associated with conventional TD learning, not zero out this egregious prediction error?

One possibility is that cortico-striatal plasticity is confined to precise temporal windows, occasioned for instance by the activity of tonically active cholinergic neurons (Bradfield, Bertran-Gonzalez, Chieng, & Balleine, 2013; Deffains & Bergman, 2015; Franklin & Frank, 2015; Lerner & Kreitzer, 2011). This window could be explicitly closed as part of the operation of control and so not lead to untoward plasticity.

There might instead be an active mechanism associated with opponency (Boureau & Dayan, 2011; Daw, Kakade, & Dayan, 2002; Lloyd & Dayan, 2019; see also Collins & Frank, 2014). That is, rather than

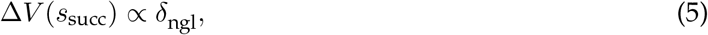

which would adjust *V* (*s*_succ_) towards 0 until *δ*_ngl_ = 0, cancelling out the reframing mechanism, we would have

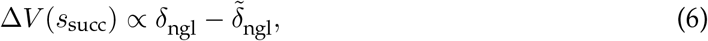

for an opponent prediction error 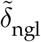. Then, Δ*V* (*s*_succ_) = 0 when

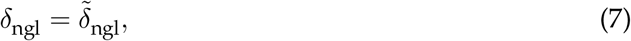

rendering reframing stable. The same argument can be made for *s*_fail_.

This perspective elucidates other cases with apparently non-zero asymptotic DA. Thus, the evidence from tasks demanding substantial work from subjects is that DA release does not inversely covary (at least strongly) with demands on vigour, but that compromising DA (e.g., by selective lesions; Salamone & Correa, 2012) compromises the willingness of subjects to overcome substantial effort costs in their active responding. If we imagine that those effort costs are conveyed by opponent terms such as 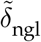, then the net influence on action in thestriatum would depend on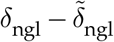, which would evidently be compromised by DA deficits.

The opponent might also help resolve a tension in our model of active avoidance between the apparent consistency of good-avoiders’ behaviour with negligible Pavlovian influence (i.e., *ω* ≈ 0) and the putative origin of positive DA on shock trials in the deployment of control. Instead of having no effect, as at present in the model, control might also be necessary for the good avoiders to overcome the malign influence of the opponent, depending on the latter’s Pavlovian effect. Of course, control might also influence the opponent (Maier & Watkins, 2005), making for a rich palette of possible interactions.

Key areas for future work include modelling the cost (Shenhav, Botvinick, & Cohen, 2013), learning (Lieder, Shenhav, Musslick, & Griffiths, 2018), and anatomical realization (putatively involving the anterior cingulate cortex; Carter, Botvinick, & Cohen, 1999) of cognitive control, along with the likely (and recursive) mesocortical DA influence over its prefrontal operation (Cools, 2019; Yee & Braver, 2018); addressing the ultimate habitization, at least for the avoidance case, of the relevant action and obviation of cognitive control (Cain, 2019); encompassing the known spatial (de Jong, Fraser, & Lammel, 2022; Menegas, Akiti, Amo, Uchida, & Watabe-Uchida, 2018) and temporal (Mohebi et al., 2019) heterogeneity in DA release; explaining the effects of pharmacological manipulation in the Go/No-Go task (Grima et al., 2022; Härmson, Grima, Panayi, Husain, & Walton, 2022); and incorporating the dorsal striatum, with its suggested focus on the instrumental components of control, and its own dopaminergicallyimpacted bias in favour of action (Collins & Frank, 2014). It will also be important to model the evolution of DA over the whole trial, broadening our current focus from DA concentrations around the cue.

In sum, we have suggested a neurocomputational architecture in which simple rules coupling action and valence are subject to a form of cognitive control whose mode of action exploits this very coupling.

## Acknowledgements

We are very grateful to Matthew Roesch and Mark Walton for their helpful comments on a previous version of the manuscript. This work was supported by the Max Planck Society and the Alexander von Humboldt Foundation.

## Methods

### 1 General model

Both tasks are modelled in essentially the same way (Fig. 3). First, an ‘internal’ decision is made when a cue arrives, at state *s*_cue_, about whether to apply self-control (ℂ= 1) or not (ℂ= 0). There then follows an ‘external’ decision about the physical action at state *s*^1^ or *s*^0^, as appropriate. It is ultimately the physical action that determines success (to *s*_succ_) or failure (to *s*_fail_) for the current trial. Following the inter-trial interval (ITI), and any additional time penalty for failure, the next trial begins.

**Figure 3:**
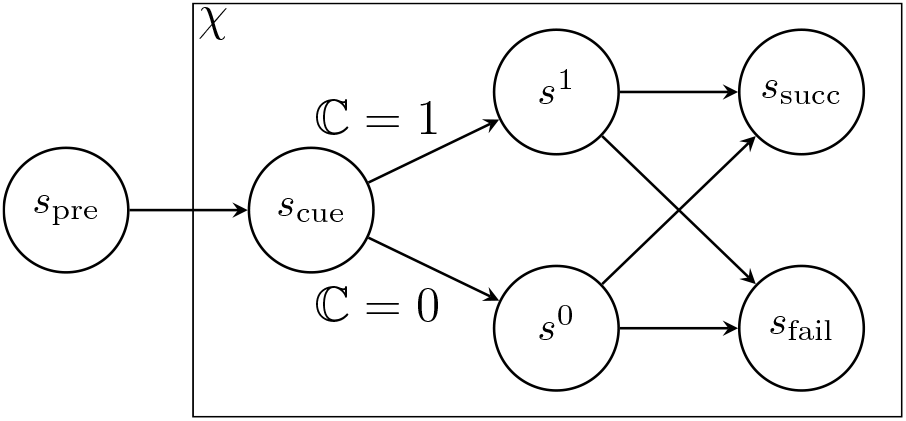
General model.

We denote trial type by *χ*. For simplicity, we assume that the initial, internal decision is made only on the trial types of most interest, i.e., on shock trials (*χ* = shk) in the mixed-valence task, and on No-Go large-reward (NGL) trials (*χ* = ngl) in the Go/No-Go task. Thus, we assume that the default choice is always ℂ = 0, only possibly deviating on these particular trial types. The choice of whether to apply self-control or not is modelled in a very simple way. For shock and NGL trials, we simply assume that there is a fixed probability *κ*_*χ*_ of applying selfcontrol,

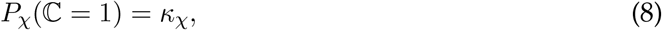

with this probability being fit to summary measures of the data as described below (Section 1.4).

As described in the main text, the importance of this internal choice is its effect on the cue-elicited temporal difference (TD) error. For ℂ = 0, we assume that *s*^0^ is in essence an extension of the cue state, and so there is really no change to the TD error elicited by cue onset, i.e.,

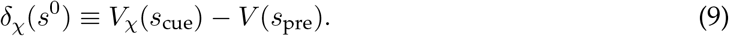

By contrast, for ℂ = 1, we assume that this is transformed to

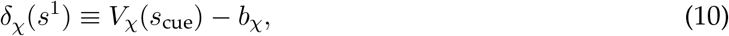

where *b*_*χ*_ is the trial-specific baseline/control signal that implements the putative reframing. As argued in the main text, we assume that *b*_*χ*_ is precisely the disutility of the shock that will be experienced on a shock trial if the animal fails to press, or is the utility of the large reward that stands to be lost on a NGL trial if the animal fails to maintain fixation.

In describing the models in greater detail below, we make use of the following common notation:

- *R*_*χ*_(*s*): immediate expected utility in current state *s* and trial type *χ*.
- *T*_*χ*_(*s*)/*T*_*χ*_(*s, a, τ*): expected time until the next state from current state *s* and trial type *χ*, potentially also conditioned on action (*a, τ*).

#### 1.1 Mixed-valence task

In this case (Gentry et al., 2016), the trial types are *χ* ∈ {rew, neu, shk}, and we assume that the animal’s external choice is between *press* and *other*, where *other* can be thought of as some alternative activity that the animal may choose to engage in (e.g., grooming) and which may itself be rewarding, but will mean that the animal fails to press on the current trial.

In the experiment of Gentry et al. (2016), there was a 5 s interval between cue onset and the insertion of the response lever; in the model, for simplicity, we assume that the external choice is made at the time of the cue, and that implementation of that choice only begins at lever insertion. Following our previous work (Lloyd & Dayan, 2019), we assume that successfully pressing on a reward trial delivers positive utility *r*_rew_ = 4, while failing to press on a shock trial leads to delivery of a punishing shock with disutility *r*_shk_ = −10. Briefly, the unpleasantness of the shock is assumed to be greater in magnitude than the pleasantness of the reward based on the (dopaminergic) evidence that neutral trials had relatively positive value for poor avoiders, in spite of an average rate of successful avoidance of around 50%; roughly speaking, for this to hold under the model, the magnitude of the disutility of shock would need to be more than twice the utility of the reward (see Lloyd & Dayan, 2019, for more detail).

If the animal chooses *press*, we assume it also chooses a latency *τ*_min_ ≤ *τ* ≤ *τ*_max_ for press completion, with *τ*_min_ = 0.5 s (for rough consistency with the data, although animals can certainly act more swiftly) and *τ*_max_ = 20 s. The latter may seem implausibly long, but also means we do not exclude the possibility that the animal chose to *press* but did not manage to complete it (e.g., due to freezing) before the *τ*_*D*_ = 10 s deadline. The (differential) value of pressing with latency *τ* is then

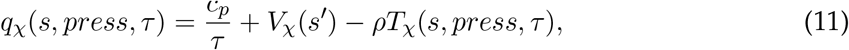

where *c*_*p*_ ≤ 0 is the assumed (hyperbolic) vigour cost associated with choosing to press at latency *τ* (cf. Niv, Daw, Joel, & Dayan, 2007); *ρ* is the average reward rate; *s* ∈ {*s*_0_, *s*_1_} (noting that the instrumental values are the same for these states);

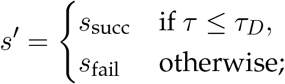

and

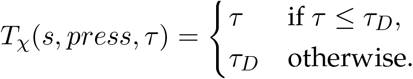

For *other*, we simply assume, again for *s* ∈ {*s*_0_, *s*_1_}, that

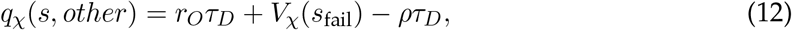

for some fixed utility rate *r*_*O*_ ≥ 0.

We assume that both instrumental and Pavlovian factors can affect speed of pressing. For presentational purposes in the main text, we refer to a single parameter *ω* that mediates the Pavlovian influence. However, in the model, we introduce a set of weights {*w*}, all *w* ≥ 0, that allow instrumental and Pavlovian factors to be balanced, but permitting Pavlovian influences on decisions about action vs. latency, and for positive vs. negative TD errors to differ (cf. Lloyd & Dayan, 2019). This does not alter the central argument, and is a simple stand-in for the complexities of how these influences are mediated. In particular, we assume the distribution of pressing latencies to follow

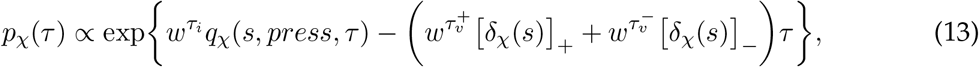

where 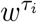 is an instrumental weight, and 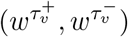 are Pavlovian weights that modulate the effect of positive and negative TD errors, respectively. This means that a positive TD error [*δ*_*χ*_(*s*)]_+_ will tend to speed up responding (since the value of a shorter *τ* will be penalized less by the term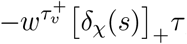, which is negative), while a negative TD error [*δ*_*χ*_ (*s*)] _−_will tend to slow responding down (since the value of a longer *τ* will be boosted more by the term

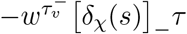, which is positive).

The over all instrumental value of pressing is then assumed to be

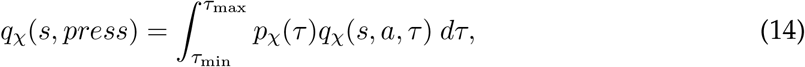

and the choice between *press* and *other* similarly assumed to be influenced by both instrumental and Pavlovian factors,

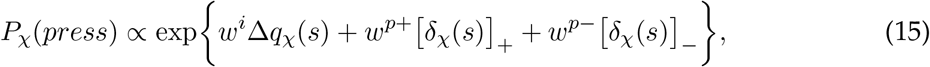

where, again, *w*^*i*^ is an instrumental weight and (*w*^*p*+^, *w*^*p*−^) are Pavlovian weights for positive and negative TD errors; and Δ*q*_*χ*_(*s*) ≡ *q*_*χ*_(*s, press*) −*q*_*χ*_(*s, other*). Thus, a positive TD error will tend to increase task engagement by boosting the probability of pressing, while a negative TD error will tend to decrease task engagement by instead boosting the probability of choosing *other*.

Note that there are two possible reasons for not pressing in the model. One is choosing *press* but failing to complete the press in time, for example because of a tendency to freeze (which we model implicitly via Eq. 13), while the other is by choosing *other*. The latter choice could be purely instrumental (i.e., the reward and effort associated with the lever press, including savings on opportunity cost of time, is not sufficiently better than the alternative) or also involve Pavlovian factors (e.g., it makes instrumental sense to press the lever, but a negative TD error promotes disengagement via a form of disappointment or frustration — via Eq. 15).

The values of success and failure states are respectively

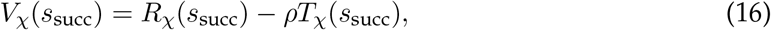

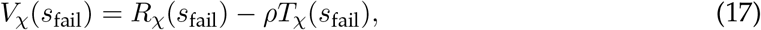

where *R*_rew_(*s*_succ_) = *r*_rew_ = 4 for a successful reward trial, and zero otherwise; *R*_shk_(*s*_fail_) = *r*_shk_ = −10 for a failed shock trial, and zero otherwise; and *T*_*χ*_(*s*_succ_) = *T*_*χ*_(*s*_fail_) = 20 s, ∀*χ*, is the ITI.

#### 1.2 Go/No-Go task

In this case (Syed et al., 2016), the task demands a slightly different choice structure. As in the mixed-valence case, there is an initial internal choice about self-control. However, we then assume that the next immediate choice facing the animal is *when to leave the nose-poke*. That is, the animal simply chooses a time *τ* to leave the nose-poke. The trial types are *χ* ∈ {gs, gl, ngs, ngl}; the value of leaving at time *τ* for both *s* ∈ {*s*^0^, *s*^1^} is assumed to be

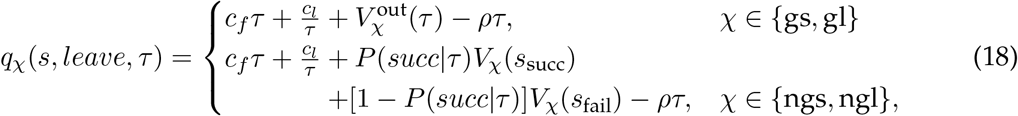

where *c*_*f*_ ≤ 0 is a cost rate assumed to be associated with maintaining fixation in the nosepoke (e.g., reflecting a decreased ability to monitor for danger); *c*_*l*_ ≤ 0 is a cost rate associated with the vigour of leaving (i.e., shorter latencies are assumed more energetically demanding); 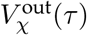 is the value on Go trials of having exited the nose-poke at time *τ* ; and *P* (*succ* |*τ*) = 𝒰 (1.7 s, 1.9 s) is the probability of success on No-Go trials of exiting the nose-poke at time *τ* (i.e., the probability that the tone has turned off before exiting — see Syed et al., 2016).

We should note here that even if the animal chooses to leave at particular time *τ* and the tone turns off *before* this time, we assume the animal will still in fact exit at time *τ* and incur the full costs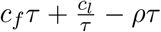. In other words, we assume that the unfolding of the animal’s leaving is non-interruptible. If we assumed that the animal could be interrupted by the tone’s turning off and pay only a fractional cost of what it actually chose (cf. Dayan, 2012), then it could make sense here to simply choose the slowest possible leaving time (which would have the lowest expected cost and, if implemented, would never result in leaving too early). Under the current formulation, the animal needs to balance, on the one hand, the vigour cost and possibility of failure if it leaves quickly and, on the other, the fixation cost and opportunity cost of time if it leaves slowly.

Just as with pressing latencies in the mixed-valence case (cf. Eq. 13), we assume that the distribution of leaving times is influenced by both instrumental and Pavlovian factors,

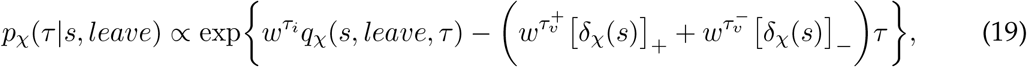

so that a positive TD error promotes leaving earlier, while a negative TD error promotes leaving later.

While the choice of leaving latency determines success or failure on No-Go trials (success is signalled by reward delivery, and failure by the turning on of the houselight — see Syed et al., 2016), we assume that on Go trials, an additional choice is required. That is, on exiting the nose-poke, the animal additionally chooses — as in the mixed-valence task — whether to subsequently *press* the lever or perform some *other* activity. Again, we can consider the value of pressing at different latencies, though this now depends on the time at which the animal exited the nose-poke; letting *t* denote the time that has elapsed since cue onset, the value of choosing to press at latency *τ* at that time (for *χ* ∈ {gs, gl}) is assumed to be

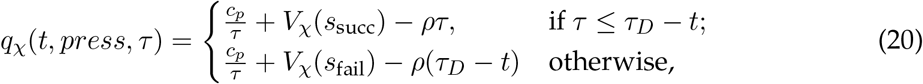

where, again, *c*_*p*_ ≤ 0 is the (vigour) cost rate associated with pressing. The instrumental value of choosing *other* is assumed to be

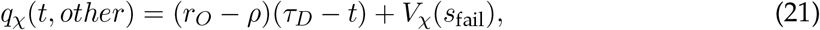

since the duration of enjoyment of this alternative activity (at reward rate *r*_*O*_ ≥ 0), and accruing opportunity cost of time (at rate *ρ*), is *τ*_*D*_ − *t* seconds.

Assuming that choice of pressing latencies given choice of *press* is governed by a softmax function,

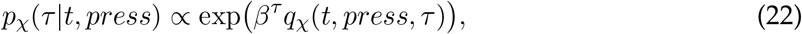

and

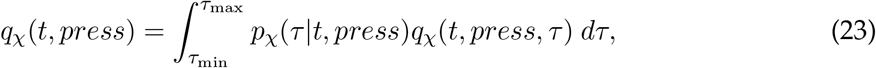

we assume that the probability of choosing *press*, as opposed to *other*, at time *t* is given by the logistic/softmax function

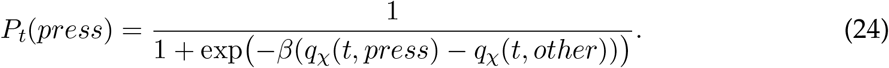

Note that we therefore assume for simplicity that the choice between *press* and *other* on exiting the nose-poke is purely instrumentally-governed. The value of having left the nose-poke at time *t* on a Go trial is then

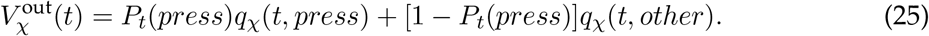

The value of success states are

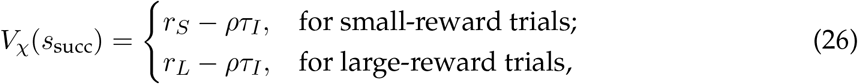

where *τ*_*I*_ = 5 s is the ITI. For fail states, we have for all trial types

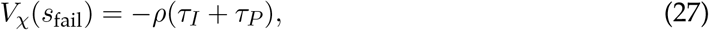

where *τ*_*P*_ = 5 s is the penalty timeout.

#### 1.3 Dopamine

As described in the main text, we assume that the Pavlovian influence is via the TD prediction error *δ*_*χ*_(*s*). While a rather large body of evidence supports the idea that this quantity is signalled by the activity of midbrain DA neurons (see citations in main text), there is also evidence that DA activity may not be its sole representational substrate. Longstanding ideas of opponency suggest that particularly in relation to punishing outcomes and their predictors, other substrates (e.g., serotonin) play a role (Boureau & Dayan, 2011; Daw et al., 2002). Indeed, one interesting feature of the results in Gentry et al. (2016) is the apparent absence of any immediate dip in DA in response to the shock cue (perhaps before control is deployed) on successful avoidance trials.

We follow Daw et al. (2002) in assuming that DA indeed only signals part of the full TD error, and principally signals transitions that are better than expected. In particular, similar (but not identical) to Daw et al. (2002), we assume that the dopaminergic component 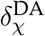 is given by

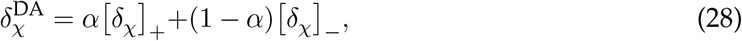

with *α* = 0.8 throughout, for simplicity.

We additionally need to consider how the quantity 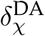, putatively represented by the firing of DA neurons, is reflected in changes in DA release measured in the accumbens (NAc). Here, we simply assume that this term is convolved with an alpha function (cf. Lloyd & Dayan, 2015),

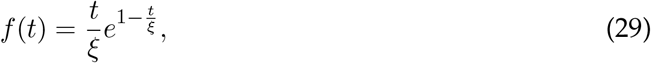

with time constant *ξ* = 0.7 s, so that changes in DA concentration relative to baseline are given by

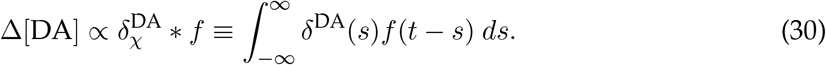

To allow for the possibility that it may take some non-trivial amount of time for control/reframing to be applied (as possibly hinted by the results of Syed et al., 2016 on successful NGL trials), for trials on which this is the case (i.e., ℂ= 1), we assume that we initially have 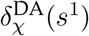 at cue onset, but that this is followed by 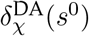 after a short, subsecond delay (we arbitrarily set this to *τ*_delay_ = 250 ms).

Note that to separately assess model-derived DA responses on success vs. failed trials, we compute the posterior probability of having employed control given success (since success and failure could occur whether or not control was employed):

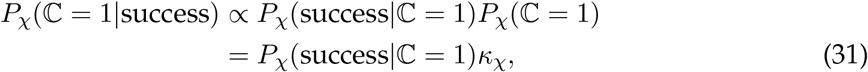

and the probability of having employed control given failure,

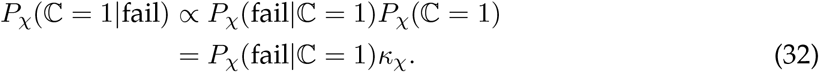

Thus, the average DA signal on successful trials is a mixture of those on which control was employed (with inferred probability *P*_*χ*_(ℂ= 1|success)) and those on which it was not (with probability *P*_*χ*_(ℂ = 0|success) = 1 − *P*_*χ*_(ℂ = 1|success)). Analogous reasoning allows for derivation of the average DA signal on failed trials.

#### 1.4 Model-fitting

For a given set of parameters **x**, the self-consistent set of differential state values and associated behaviour can be found using value iteration (Mahadevan, 1996). For each task, we fitted parameters (see Tables 1 and 2) to minimize the difference between animal and model behaviour. In particular, we defined an error function

**Table 1:** Model parameters used for mixed-valence task and best-fitting free parameter values for goodand poor-avoiders.

| model parameters |  |  |  |
| --- | --- | --- | --- |
| fixed | free | good | poor |
| $r_{rew} = 4$ | $w^i$ | 2.45 | 0.01 |
| $r_{neu} = 0$ | $w^{p+}$ | 0.00 | 0.82 |
| $r_{shk} = -10$ | $w^{p-}$ | 0.01 | 0.74 |
| $r_O = 0$ | $w^{\tau_i}$ | 41.28 | 0.00 |
| $c_p = -0.1$ | $w^{\tau_p^+}$ | 0.97 | 1.06 |
| $b_{shk} = r_{shk}$ | $w^{\tau_p^-}$ | 0.00 | 0.00 |
| $\alpha = 0.8$ | $\kappa$ | 0.36 | 0.50 |
| $\tau_{delay} = 250$ ms | | | |

**Table 2:**
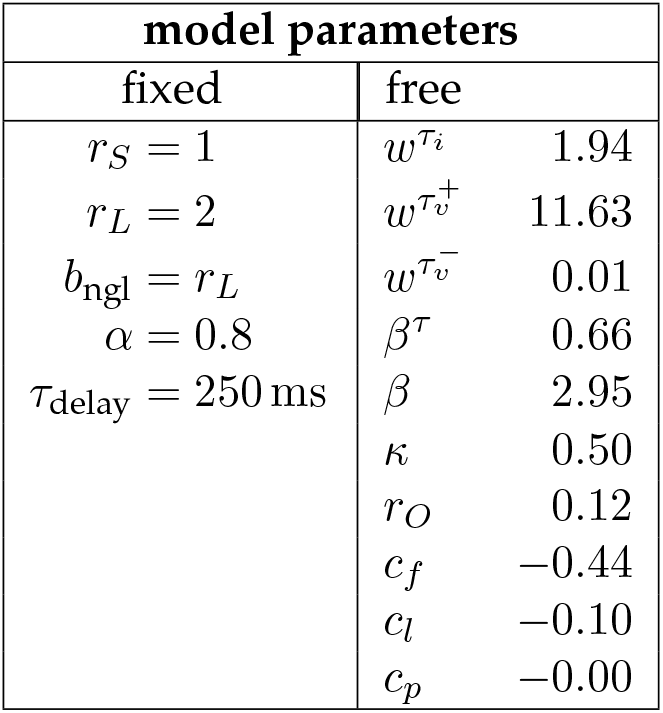
Model parameters used for Go/No-Go task and best-fitting free parameter values.

| model parameters |  |  |
| --- | --- | --- |
| fixed | free |  |
| $r_S = 1$ | $w^{\tau_i}$ | 1.94 |
| $r_L = 2$ | $w^{\tau_v^+}$ | 11.63 |
| $b_{ngl} = r_L$ | $w^{\tau_v^-}$ | 0.01 |
| $\alpha = 0.8$ | $\beta^\tau$ | 0.66 |
| $\tau_{delay} = 250$ ms | $\beta$ | 2.95 |
| | $\kappa$ | 0.50 |
| | $r_O$ | 0.12 |
| | $c_f$ | -0.44 |
| | $c_l$ | -0.10 |
| | $c_p$ | -0.00 |

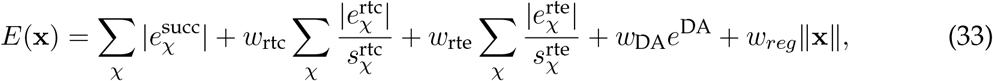

Where 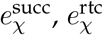, 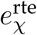, and 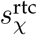are the differences between the average performance of the animals and the model (for expected success rates, reaction times for correct trials, and reaction times for error trials, respectively); 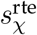 and 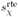 are the standard errors from the experimental data (so that the larger the standard error, the less importance we place on precise fitting of that measurement); *w*_rtc_ ≥ 0 and *w*_rte_ ≥ 0 are weights that determine the relative importance we give to fitting the reaction times compared to the success rates (in practice, set to these to the same value: for the mixed-valence task, we fixed *w*_rtc_ = *w*_rte_ = 0.25; for Go/No-Go, we fixed *w*_rtc_ = *w*_rte_ = 0.018); and *w*_*reg*_ is a (L1) regularization weight that determines how strongly we wish to discourage large values in the parameters **x** (we set *w*_*reg*_ = 0.01 when fitting good avoidance in the mixed-valence task, since the instrumental weights can grow arbitrarily large in this case, and otherwise set it to zero). The term *e*^DA^ is an error term that we used only when fitting data from the mixed-valence task, and is specifically the absolute difference between peak DA for food and shock trials produced by the model — this was to better reproduce the striking similarity between food- and shock-trial DA transients observed in the data (we set *w*_DA_ = 0.1 for both good and poor avoiders).

This latter point deserves amplification. For the good avoiders, behaviour is consistent with a purely instrumental policy, with Pavlovian weights ≈ 0 (see Table 1). This implies that behaviour in the model does not vary with *δ*_*χ*_ (or therefore the value of *κ*) — the model reliably presses on shock trials in any case. *κ* is only identifiable when additionally considering how the model’s implied DA pattern compares with the data, hence the fitting weight *w*_DA_ above. Why though would control be at all necessary if behaviour were purely instrumental, since there would then apparently be no danger of Pavlovian misbehaviour? As mentioned in the main text, one possibility is that an opponent signal might also influence behaviour, which controlled dopamine release would be required to overcome. Indeed, note that good-avoiders still freeze in response to shock cues (cf. Figure 4i of Gentry et al., 2016). We leave these subtleties to future work.

## 2 Data analysis

The data from Syed et al. (2016) were downloaded from https://data.mrc.ox.ac.uk. Following Syed et al. (2016), data were smoothed using a 0.5 s moving window and baselined by subtracting the average signal during the 0.5 s period before cue onset. As a basic test of the hypothesis that the cue-evoked DA response would be greater on failed No-Go large-reward (NGL) trials than on successful NGL trials, we integrated the DA signal over the 1 s period immediately following cue-onset, averaged this measure for each session, and applied a (paired, one-tailed) *t*-test to these session-wise averages.

## Supplementary figures

**Figure S1:**
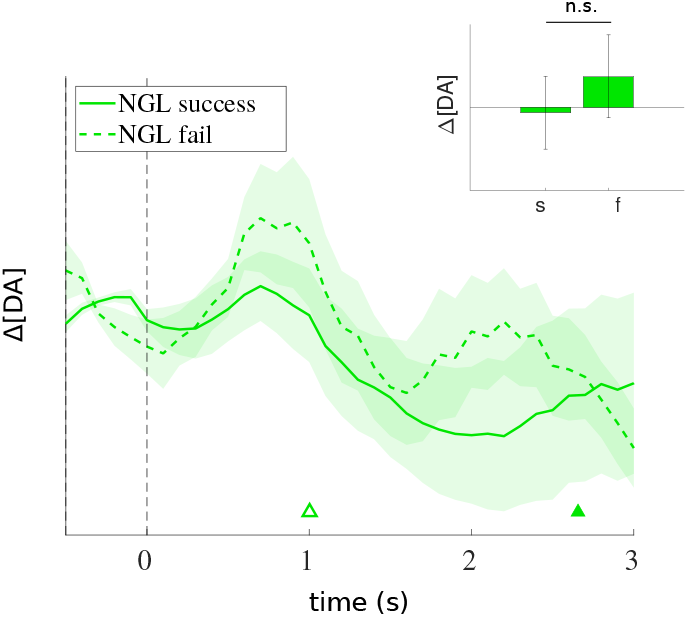
Average cue-aligned (at 0 s) change in DA release (±SEM) on successful vs. failed No-Go large-reward (NGL) trials in Syed et al. (2016); triangles indicate average RT for success (filled) vs. failure (unfilled) trials. Inset: average integrated DA signal (±SEM) over the first 1 s following cue-onset on successful (s) vs. failed (f) trials; *t*(8) = 1.04,, *p* = .16 (n.s.), one-tailed.

## References

Bennett, D., Davidson, G., & Niv, Y. (2022). A model of mood as integrated advantage. Psychological Review, 129(3), 513–541.

Berridge, K. (2007). The debate over dopamine’s role in reward: the case for incentive salience. Psychopharmacology, 191, 391–431.

Boureau, Y.-L., & Dayan, P. (2011). Opponency revisited: Competition and cooperation between dopamine and serotonin. Neuropsychopharmacology, 36(1), 74–97.

Bradfield, L., Bertran-Gonzalez, J., Chieng, B., & Balleine, B. (2013). The thalamostriatal pathway and cholinergic control of goal-directed action: Interlacing new with existing learning in the striatum. Neuron, 79(1), 153–166.

Brischoux, F., Chakraborty, S., Brierley, D., & Ungless, M. (2009). Phasic excitation of dopamine neurons in ventral VTA by noxious stimuli. Proceedings of the National Academy of Sciences of the United States of America, 106(12), 4894–4899.

Budygin, E., Park, J., Bass, C., Grinevich, V., Bonin, K., & Wightman, R. (2012). Aversive stimulus differentially triggers subsecond dopamine release in reward regions. Neuroscience, 201, 331–337.

Cain, C. K. (2019). Avoidance problems reconsidered. Current Opinion in Behavioral Sciences, 26, 9–17.

Carter, C. S., Botvinick, M. M., & Cohen, J. D. (1999). The contribution of the anterior cingulate cortex to executive processes in cognition. Reviews in the Neurosciences, 10(1), 49–58.

Collins, A., & Frank, M. (2014). Opponent actor learning (opal): modeling interactive effects of striatal dopamine on reinforcement learning and choice incentive. Psychological Review, 121(3), 337–366.

Cools, R. (2019). Chemistry of the adaptive mind: Lessions from dopamine. Neuron, 104, 113–131.

Crockett, M., Clark, L., & Robbins, T. (2009). Reconciling the role of serotonin in behavioral inhibition and aversion: Acute tryptophan depletion abolishes punishment-induced inhibition in humans. Journal of Neuroscience, 29(38), 11993–11999.

Daw, N., Kakade, S., & Dayan, P. (2002). Opponent interactions between serotonin and dopamine. Neural Networks, 15, 603–616.

Dayan, P. (2012). Instrumental vigour in punishment and reward. European Journal of Neuroscience, 35(7), 1152–1168.

Dayan, P., Niv, Y., Seymour, B., & Daw, N. D. (2006, October). The misbehavior of value and the discipline of the will. Neural Networks, 19(8), 1153–60.

Deffains, M., & Bergman, H. (2015). Striatal cholinergic interneurons and cortico-striatal synaptic plasticity in health and disease. Movement Disorders, 30(8), 1014–1025.

de Jong, J. W., Afjei, S. A., Dorocic, I. P., Peck, J. R., Liu, C., Kim, C. K., … Lammel, S. (2019). A neural circuit mechanism for encoding aversive stimuli in the mesolimbic dopamine system. Neuron, 101(1), 133–151.

de Jong, J. W., Fraser, K. M., & Lammel, S. (2022). Mesoaccumbal dopamine heterogeneity: What do dopamine firing and release have to do with it? Annual Review of Neuroscience, 45, 109–129.

Franklin, N. T., & Frank, M. J. (2015). A cholinergic feedback circuit to regulate striatal population uncertainty and optimize reinforcement learning. Elife, 4, e12029.

Gentry, R., Lee, B., & Roesch, M. (2016). Phasic dopamine release in the rat nucleus accumbens predicts approach and avoidance performance. Nature Communications, 7(131154).

Grima, L., Panayi, M., Härmson, O., Syed, E., Manohar, S., Husain, M., & Walton, M. (2022). Nucleus accumbens D1-receptors regulate and focus transitions to reward-seeking action. Neuropsychopharmacology.

Guitart-Masip, M., Beierholm, U., Dolan, R., Duzel, E., & Dayan, P. (2011). Vigor in the face of fluctuating rates of reward: an experimental examination. Journal of Cognitive Neuroscience, 23(12), 3933–3938.

Härmson, O., Grima, L., Panayi, M., Husain, M., & Walton, M. (2022). 5-HT2C receptor perturbation has bidirectional influence over vigour and restraint. Psychopharmacology, 239, 123–140.

Huys, Q., Cools, R., Gölzer, M., Friedel, E., Heinz, A., Dolan, R., & Dayan, P. (2011). Disentangling the roles of approach, activation and valence in instrumental and pavlovian responding. PLoS Computational Biology, 7(4), e1002028.

Kim, H. R., Malik, A. N., Mikhael, J. G., Bech, P., Tsutsui-Kimura, I., Sun, F., … Uchida, N. (2020). A unified framework for dopamine signals across timescales. Cell, 183(6), 1600– 1616.

Lerner, T., & Kreitzer, A. (2011). Neuromodulatory control of striatal plasticity and behavior. Current Opinion in Neurobiology, 21(2), 322–327.

Lieder, F., Shenhav, A., Musslick, S., & Griffiths, T. L. (2018). Rational metareasoning and the plasticity of cognitive control. PLoS computational biology, 14(4), e1006043.

Lloyd, K., & Dayan, P. (2015). Tamping ramping: Algorithmic, implementational, and computational explanations of phasic dopamine signals in the accumbens. PLoS Computational Biology, 11(12), e1004622.

Lloyd, K., & Dayan, P. (2019). Pavlovian-instrumental interactions in active avoidance: The bark of neutral trials. Brain research, 1713, 52–61.

Mahadevan, S. (1996). Average reward reinforcement learning: Foundations, algorithms, and empirical results. Machine Learning, 22, 159–196.

Maier, S. F., & Watkins, L. R. (2005). Stressor controllability and learned helplessness: the roles of the dorsal raphe nucleus, serotonin, and corticotropin-releasing factor. Neuroscience & Biobehavioral Reviews, 29(4-5), 829–841.

McClure, S., Daw, N., & Montague, P. (2003). A computational substrate for incentive salience. Trends in Neurosciences, 26(8), 423–428.

Menegas, W., Akiti, K., Amo, R., Uchida, N., & Watabe-Uchida, M. (2018). Dopamine neurons projecting to the posterior striatum reinforce avoidance of threatening stimuli. Nature neuroscience, 21(10), 1421–1430.

Mohebi, A., Pettibone, J. R., Hamid, A. A., Wong, J.-M. T., Vinson, L. T., Patriarchi, T., … Berke, J. D. (2019). Dissociable dopamine dynamics for learning and motivation. Nature, 570(7759), 65–70.

Montague, P., Dayan, P., & Sejnowski, T. (1996). A framework for mesencephalic dopamine systems based on predictive hebbian learning. The Journal of Neuroscience, 16(5), 1936– 1947.

Niv, Y., Daw, N., Joel, D., & Dayan, P. (2007). Tonic dopamine: opportunity costs and the control of response vigor. Psychopharmacology, 191(3), 507–520.

Oleson, E., & Cheer, J. (2013). On the role of subsecond dopamine release in conditioned avoidance. Frontiers in Neuroscience, 7, 101–109.

Oleson, E., Gentry, R., Chioma, V., & Cheer, J. (2012). Subsecond dopamine release in the nucleus accumbens predicts conditioned punishment and its successful avoidance. The Journal of Neuroscience, 32(42), 14804–14808.

Salamone, J., & Correa, M. (2012). The mysterious motivational functions of mesolimbic dopamine. Neuron, 76, 470–485.

Schultz, W., Dayan, P., & Montague, P. (1997). A neural substrate of prediction and reward. Science, 275, 1593–1599.

Shenhav, A., Botvinick, M., & Cohen, J. (2013). The expected value of control: An integrative theory of anterior cingulate function. Neuron, 79, 217–240.

Starkweather, C. K., & Uchida, N. (2021). Dopamine signals as temporal difference errors: recent advances. Current Opinion in Neurobiology, 67, 95–105.

Sutton, R. (1988). Learning to predict by the methods of temporal differences. Machine Learning, 3(1), 9–44.

Swart, J., Froböse, M., Cook, J., Geurts, D., Frank, M., Cools, R., & den Ouden, H. (2017). Catecholaminergic challenge uncovers distinct pavlovian and instrumental mechanisms of motivated (in)action. eLife, 6(e22169).

Syed, E., Grima, L., Magill, P., Bogacz, R., Brown, P., & Walton, M. (2016). Action initiation shapes mesolimbic dopamine encoding of future rewards. Nature Neuroscience, 19(1), 34–36.

Wenzel, J., Oleson, E., Gove, W., Cole, A., Gyawali, U., Dantrassy, H., … Cheer, J. (2018). Phasic dopaminergic signals in the nucleus accumbens that cause active avoidance require endocannibinoid mobilization in the midbrain. Current Biology, 28(9), 1392–1404.

Yee, D. M., & Braver, T. S. (2018). Interactions of motivation and cognitive control. Current opinion in behavioral sciences, 19, 83–90.

Young, A. (2004). Increased extracellular dopamine in nucleus accumbens in response to unconditioned and conditioned aversive stimuli: studies using 1 min microdialysis in rats. Journal of Neuroscience Methods, 138(1), 57–63.

